# Tattoo ink induces inflammation in the draining lymph node and impairs the immune response against a COVID-19 vaccine

**DOI:** 10.1101/2024.12.18.629172

**Authors:** Arianna Capucetti, Juliana Falivene, Chiara Pizzichetti, Irene Latino, Luca Mazzucchelli, Vivien Schacht, Urs Hauri, Andrea Raimondi, Tommaso Virgilio, Alain Pulfer, Simone Mosole, Llorenec Grau-Roma, Wolfgang Bäumler, Martin Palus, Louis Renner, Daniel Ruzek, Gabrielle Goldman Levy, Milena Foerster, Kamil Chahine, Santiago F. Gonzalez

**Author notes:** Equal contribution (Arianna Capucetti, Juliana Falivene). **Corresponding Author:** Santiago Fernandez Gonzalez, Institute for Research in Biomedicine, via Francesco Chiesa 5. CH-6500 Bellinzona. Switzerland.

## Abstract

Despite safety concerns regarding the toxicity of tattoo ink, no studies have reported the consequences of tattooing on the immune response. In this work, we have characterized the transport and accumulation of different tattoo inks in the lymphatic system using a murine model. Upon quick lymphatic drainage, we observed that macrophages mainly capture the ink in the lymph node (LN). An initial inflammatory reaction at local and systemic levels follows ink capture. Notably, the inflammatory process is maintained over time as we observed clear signs of inflammation in the draining LN two months following tattooing. In addition, the capture of ink by macrophages was associated with the induction of apoptosis in both human and murine models. Furthermore, the ink accumulated in the LN altered the immune response against a COVID-19 vaccine. We observed a reduced antibody response following vaccination with a mRNA-based SARS-CoV-2 vaccine, which was associated with a decreased expression of the Spike protein in macrophages in the draining LN. Considering the unstoppable trend of tattooing in the population, our results are crucial in informing the toxicology programs, policymakers, and the general public regarding the potential risk of the tattooing practice associated with an altered immune response.

## Introduction

Tattooing has become widespread, particularly for young people. According to the latest studies, it is estimated that globally, almost one out of five individuals have tattoos^1^, with the USA having the highest prevalence with more than 30% of tattooed individuals^2,3^. During the tattooing process, repeated penetration by needles into the dermal layer of the skin introduces different types of pigments following a pattern. The permanence of tattoos is achieved by using pigments that are not readily soluble in bodily fluids, often formed by a complex mixture of pigment binders, solvents, and additives^4^. While carbon black is the primary pigment used in black tattoos, colored tattoos typically contain industrial organic pigments originally intended for plastics, varnishes, or paints^5,6^.

Despite the increasing prevalence of tattooing, the regulation of tattoo inks is less stringent than that of the pharmaceutical industry or other products intended for human use. Even though toxicological data may be available for some ink ingredients^7^, studies on *in vivo* interactions of the ink components and their fate within the body are rare. The lack of comprehensive regulations has prompted the identification of risks associated with tattooing and the development of specific information to ensure that individuals can make informed decisions before undergoing this procedure^8,9^. In Europe, the composition of the ink has been regulated since 2022 by the REACH program, which intends to harmonize the legislation through the member states^10^. However, the potential health implications of this form of body art have become a subject of increasing concern^5^.

Previous studies have shown that macrophages are the primary cells responsible for ink uptake in the skin^11^. In conjunction with tattooed skin, pigmented and enlarged lymph nodes (LN) and several adverse immune-related reactions have been reported in tattooed individuals for decades^12–18^. Notably, studies performed in mouse models have shown the accumulation of tattoo ink in the LN draining the tattoo site, mimicking what has been observed in humans and primates^19,20^. Despite the central role of the LN in the development of appropriate immune responses, the mechanisms and potential consequences behind the immunological reactions upon tattooing remain unknown. Moreover, many concerns regarding complications from vaccination and other medical procedures in tattooed individuals have been raised over the years, especially during the mass vaccination against SARS-CoV-2. Although is recommended to avoid vaccination on fresh tattoos, no specific studies have evaluated the impact of tattoos on vaccination efficacy or the risk of infections associated with tattoos^21^.

In summary, this study elucidates the transport and accumulation of tattoo inks in the draining LN, characterizing the inflammatory response to tattooing and evaluating how tattooing affects the humoral immune response to a COVID-19 vaccine. Our work provides a comprehensive analysis of the immunotoxicology of tattooing.

## Results

### The lymphatic system transports tattoo ink, which accumulates in the synodical areas of the draining LN

We selected three of the most used colors of ink, black, red, and green, from one of the major world providers (Intenze) and confirmed that the composition of the inks was concordant with the REACH regulation^10^ (**Supp.** **Table 1**). Then, we set up a mouse model, tattooing an area of around 25 mm^2^ in the footpad of the mice (**Fig. 1A, E upper panels**). Next, we confirmed by electron microscopy the presence of ink in both the epidermis and the dermis of the skin, associated with cells resembling keratinocytes, macrophages, and the collagen matrix (**Supp. Fig. 1A**). To evaluate the transport of unretained ink from the skin area to the draining LN, we surgically exposed the mouse hindlimb^22^ and we performed intravital imaging following footpad tattooing (**Fig. 1B**). We observed that the ink disseminates exclusively via popliteal lymphatic vessels, reaching an initial peak at 10 min following tattooing (**Fig. 1C, D**).

**Fig. 1.**
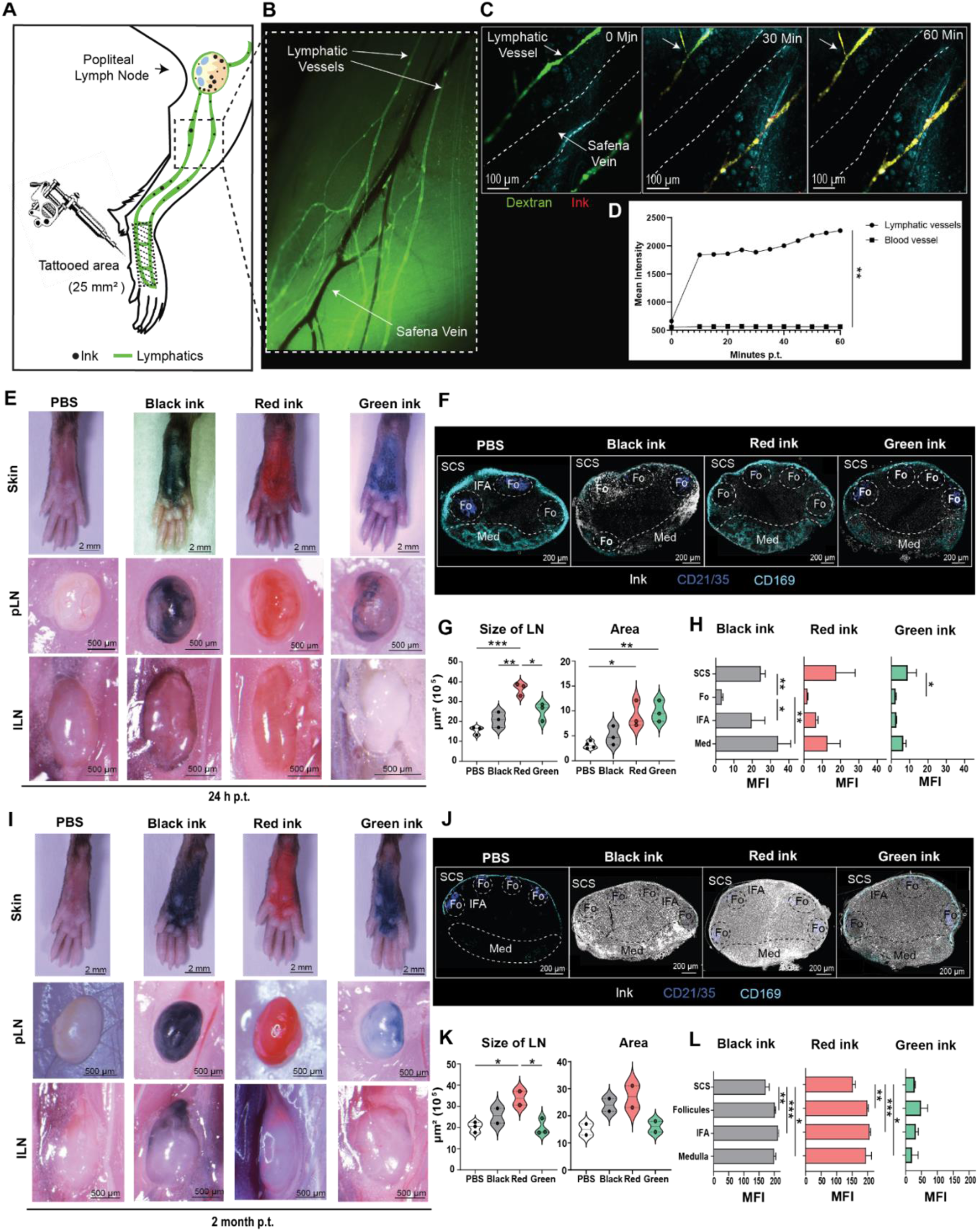
Tattoo ink is transported by the lymphatic system and accumulates in the draining lymph node. **(A)** Graphic representation of the lymphatic drainage in the tattoo murine model employed. **(B)** Fluorescence micrograph showing the imaged area located in the calf of the mouse. **(C)** Representative intravital 2-photon micrographs acquired at 0-, 30-, and 60-min post-tattooing with red ink. **(D)** Quantification of the mean intensity of the red ink autofluorescence inside the lymphatic and the blood vessels over 60 min following tattooing. Pearson correlation analysis (r=0.7690, p (two-tailed)=0.0035) **(E)** Snapshots showing the tattooed area on the mouse footpad and the ink accumulation in the draining LNs at 24 h or two months **(I)** after tattooing with commercial True Black, Bright Red, or Pure Green inks. Representative confocal micrographs following the ink distribution in the different areas of the LN at 24 h p.t. **(F)** and two months p.t **(J)**: SCS stands for subcapsular sinus; Fo, B cell follicle; Med, medullary region; IFA, interfollicular area. **(G)** Violin plot representing the average size of the pLN and the area occupied by ink signal at 24 h **(K)** and two months p.t. **(H)** Quantification of the MFI of the inks in the different regions of the LN at 24 h **(G-H)** and two months p.t. **(K-L)** In **G-H** and **K-L,** one out of two or more independent experiments is shown; Data are presented as mean ± SD or Median and 25th (bottom) and 75th (top) percentiles. One-way analyses of variance (ANOVA) followed by Bonferroni correction for multiple comparisons. Differences between groups were considered significant with p-value < 0.05 (* p < 0.05, ** p < 0.01, *** p < 0.001).

Furthermore, the transport of ink via the lymphatic system was associated with a prominent accumulation in the popliteal LN (pLN) observed at 24 h post-tattooing (p.t.) for all the tested inks (**Fig. 1E, middle panels**). Interestingly, we also observed the presence of ink in the lumbar LN with two of the tested inks (black and red), suggesting that unretained ink disseminates further via the efferent lymphatics (**Fig. 1E, lower panels**). Next, we performed confocal microscopy to determine the pLN areas where the ink pigments accumulate (**Fig. 1F).** In addition, we measured the mean fluorescent intensity (MFI) associated with the ink, observing that at 24 h p.t., there was a significant increase in the size of the pLN and in the area occupied by the ink, compared to the PBS-tattooed control (**Fig. 1G**). Moreover, at this point, the pigment signal mainly accumulated in the subcapsular and medullary areas of the LN (**Fig. 1H**). To evaluate the dynamics of ink pigment accumulation over extended periods, we examined the presence of pigments at two months p.t., detecting an increase in the signal in both the popliteal and lumbar LNs (**Fig. 1I-K, Supp. Fig. 1B**). Furthermore, we observed that ink was also present in the paracortical area of the LN (**Supp. Fig. 1B**). Remarkably, we observed a similar distribution in all the human samples from LN biopsies evaluated collected from patients who had a tattoo in the LN-drained area at least several months before the biopsy (**Supp. Fig. 1D**).

### The capture of ink in the LN is mainly associated with medullary macrophages

To evaluate the cell type responsible for the capture of the ink particles in the sinusoidal areas of the pLN we performed confocal staining of the main phagocytic populations: subcapsular sinus macrophages (SSM; CD169^+^, CD11c^dim^, F4/80^-^), medullary macrophages (MM; CD169^+^, CD11c^-^, F4/80^+^) and dendritic cells (DC, CD169^-^, CD11c^+^, F4/80^-^) at 24 h and two months p.t. with black (**Fig. 2A, B**), red (**Fig. 2C, D**) and green ink (**Fig. 2E, F**). Quantification of the images demonstrated that MM are the primary cell type responsible for the ink particle uptake, compared to the other two cell types analyzed at an initial stage and at two months p.t. (**Fig. 2B, 2D, 2F**). Furthermore, we confirmed the identity of these cells by using a transgenic mouse model expressing the CX3CR1 (**Supp. Fig 2A**). Additionally, we observed that DC can capture some ink. However, most ink particles were associated with macrophages (**Fig. 2B, 2D, 2F**; **Supp. Fig. 2B**). Next, we performed resin embedding of the pLN (**Fig. 2G, H**) and analyzed the samples using transmission electron microscopy. We confirmed that most ink particles were associated with phagocytic cells displaying a macrophage-like morphology at 24 h p.t. (**Fig. 2 H, I**). Moreover, we could observe multiple phagocytic vacuoles containing electrodense material (**Fig. 2I**). When we evaluated the structure of the same cells at two months p.t., we observed an increase in the size of the cells and the amount of ink associated with macrophages in the medullary region and the formation of giant cells with multiple phagocytic vacuoles containing ink particles (**Fig. 2K, L**). Interestingly, we confirmed that tattoo ink particles are also associated with CD68^+^ or CD163^+^ macrophages in two biopsies of LNs from tattooed patients (**Supp. Fig. 2C** and **2F**, respectively) and observed numerous giant cells containing ink particles deposited, showing a similar morphology to the ones described in the murine model (**Supp. Fig. 2E**).

**Fig. 2.**
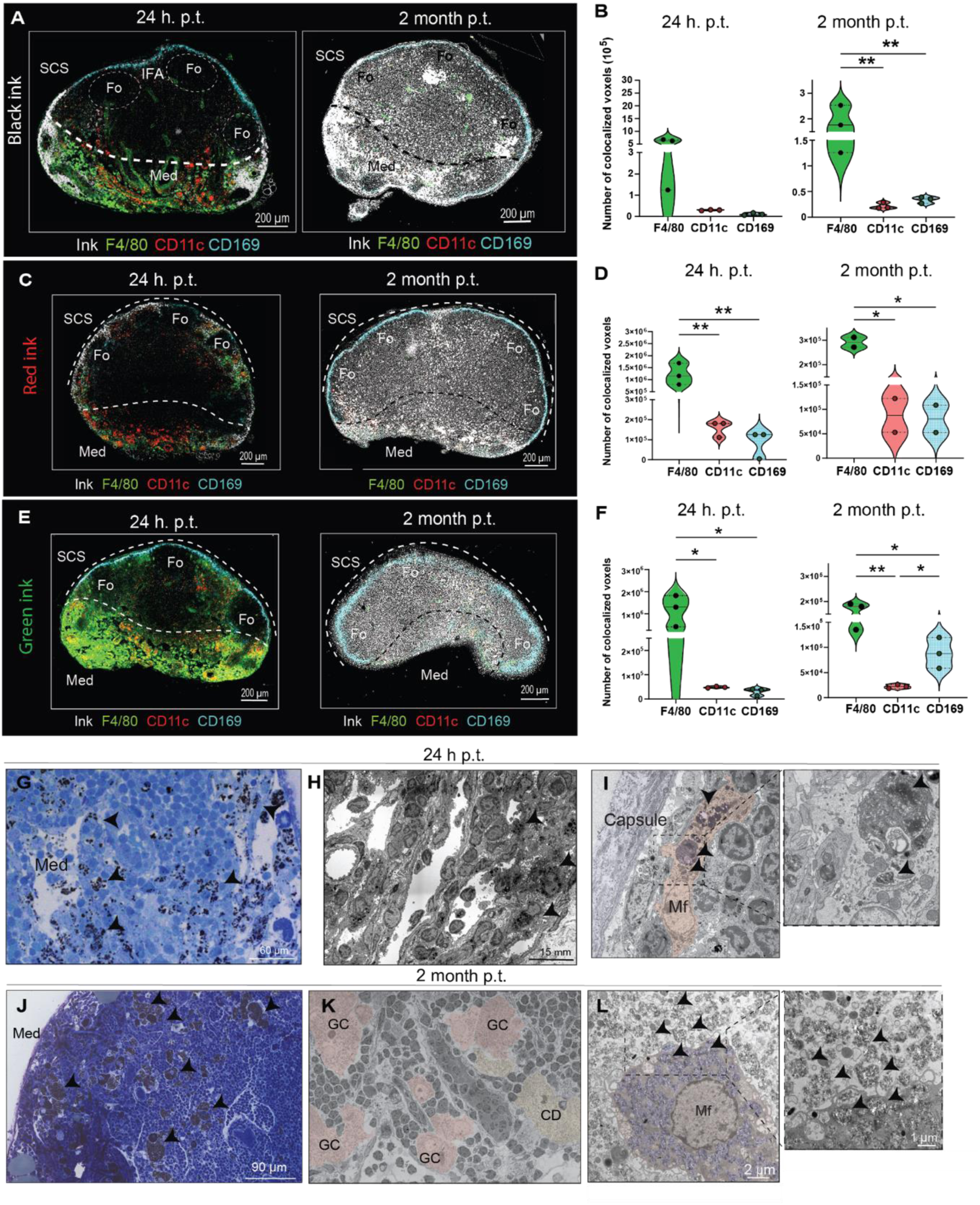
Medullary macrophages capture and retain tattoo ink. **A)** Representative confocal micrographs showing the colocalization of black, (**C**) red, and (**E**) green ink with the main phagocytic populations in the LN at 24 h (left panels) and two months (right panels) p.t. (**B**) Violin plot representing the quantification of colocalized voxels of black, (**D**) red, and (**F**) green ink with the major phagocytic populations in the draining LN at 24h (left) and two months (right) p.t..**(G)** Toluidine Blue staining of the pLN at 24 h and two months p.t. (**J**). Black arrowheads indicate the presence of ink. (**H**) Representative EM micrographs of the medullary sinuses at 24 h and two months p.t. (**K**), showing the presence of macrophages with multiple ink-containing vacuoles at 24h (**I**) and two months p.t. (**L**). Giant cell (GC); Cell debris (CD). One out of two or more independent experiments is shown. Data are presented as median and 25th (bottom) and 75th (top) percentiles. One-way ANOVA followed by Bonferroni correction for multiple comparisons (* p < 0.05, ** p < 0.01, *** p < 0.001).

### Tattoo ink induces the death of macrophages

To study the immune cell dynamics in the draining LN following the arrival of tattoo ink, we measured the total number of cells in the pLN of mice by FACS during the first five days following tattooing. We observed a significant increase in the total number of macrophages during the first six h p.t., with the red and black ink, which were followed by a significant decrease at 12 h and 24 h p.t., respectively (**Fig. 3A, Supp. Fig. 3A** right). Regarding the green ink, macrophage decrease was only observed at 120 h p.t. (**Supp. Fig. 3A** left). Thus, we hypothesized that the observed disappearance might be associated with the toxicity of the unretained ink particles in the pLN. Furthermore, we observed by EM that some of the ink-associated macrophages in the medullary sinuses of the pLN seem to undergo morphological changes such as the loss of plasma membrane integrity (**Fig. 3 B, left**) or the formation of blebs (**Fig. 3B, right**), characteristic of cell death stages^25^. Therefore, to better characterize the mechanism of ink-associated cell death, we isolated mesenchymal stem cells from the bone marrow of mice, differentiated them into macrophages, and added different concentrations of the three inks (**Supp. Fig. 3B**). Firstly, we confirm a concentration-dependent uptake of all the inks by *in vitro*-cultured macrophages that resembles the one observed in the *in vivo* studies (**Fig. 3C, Supp. Fig. 3C**). Then, to confirm the presence of ink-induced cell death we cultured the ink-treated macrophages in the presence of propidium iodide (PI), a marker of necrosis, and quantified the number of PI^+^ cells using imageXpress^®^ (**Fig. 3D, E**). Following this approach, we observed a significant increase in the number of PI^+^ cells at 12 h post ink addition (p.i.a) with the red and the black inks compared to the control group. However, we could not see any toxic effect in the samples treated with green ink (**Fig. 3E**). Moreover, we confirmed by flow cytometric analysis that all the tested inks induced different levels of apoptosis (Annexin V^+^, PI^-^ cells) in the macrophages during the first 48 h p.i.a. Indeed, the addition of red ink was followed by a significant 7-fold induction of apoptosis at 24 h, while black and green inks induced significant levels of apoptosis at 48 h p.i.a. (**Fig. 3F**). Importantly, we observed that macrophages differentiated from human peripheral blood mononuclear cells (PBMC) (**Supp. Fig. 3D**), showed similar trends, in terms of ink capture and ink-induced cell death, when cultured with all the selected inks (**Supp. Fig. 3E-H**). However, some differences were observed regarding the time of death induction. Both black and green inks induced significant levels of apoptosis already at 24 h p.i.a. (**Supp. Fig. 3F** and **3H**, respectively), while the red ink induced necrosis at 24 h, followed by apoptosis at 48 h p.i.a (**Supp. Fig. 3G**).

**Fig. 3.**
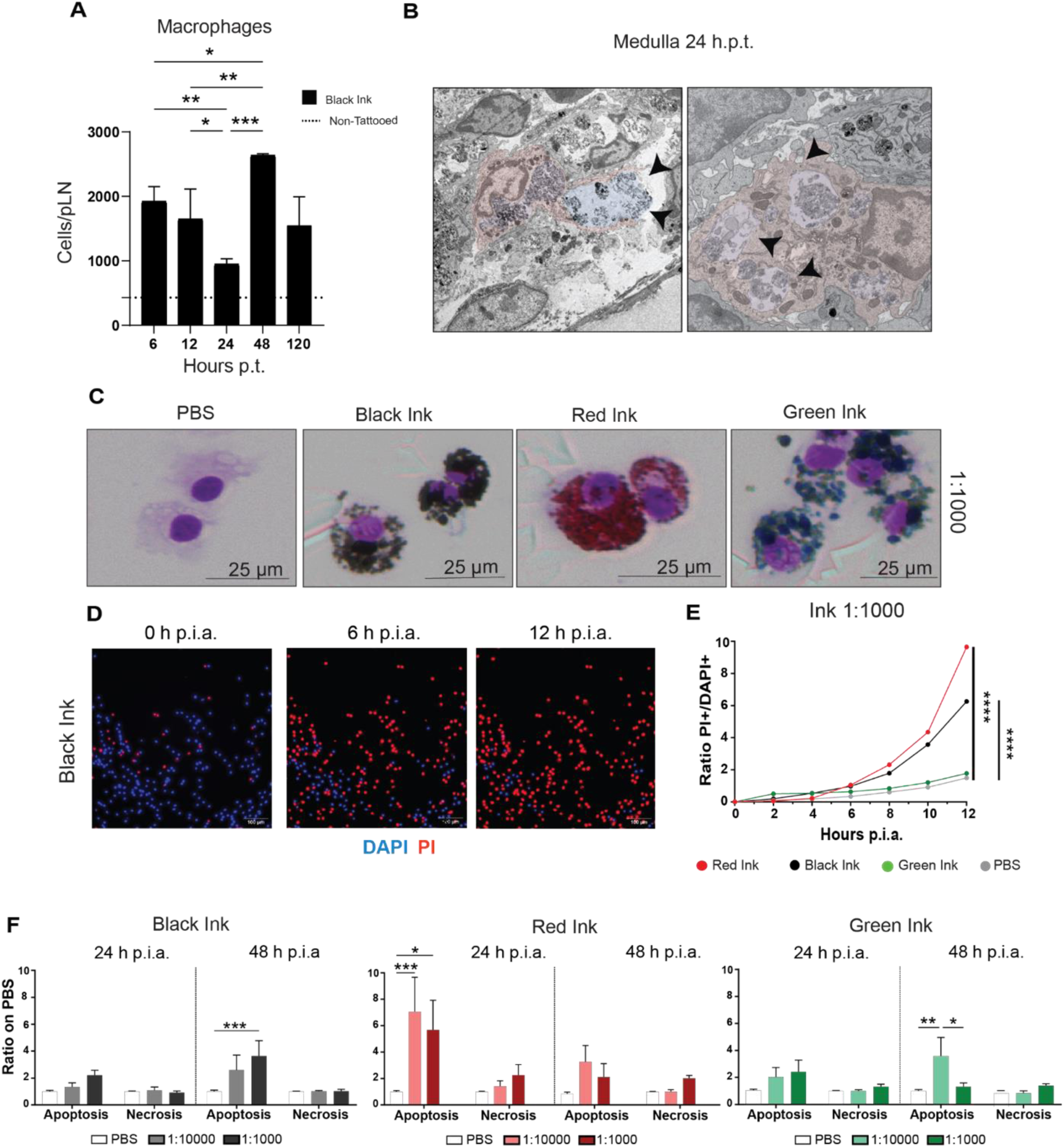
Tattoo ink induces macrophage death. (**A**) Absolute number of macrophages in the pLN of tattooed animals during the 120 h following footpad tattooing with black ink. n=4 representative of two independent experiments. (**B**) Representative EM micrographs of ink-captured macrophage-like cells (fake colored in red). Black arrowheads show signs of distress, such as the loss of plasma membrane integrity and the formation of blebs. (**C**) Representative cytospin images showing the capture of black, red, and green ink (ink dilution 1:1000) by bone marrow-derived macrophages. (**D**) Representative snapshots of DAPI and PI stained macrophages at 6 and 12 h p.i.a. and quantification plot (**E**) showing the ratio of PI^+^ on DAPI^+^ cells after adding inks. Pearson correlation analysis of black vs PBS groups (r=0.9933, p (two-tailed)<0.0001), red vs PBS groups (r=0.9817, p (two-tailed)<0.0001), and green vs PBS groups (r=0.9739, p (two-tailed)=0.0002). (**F**) Flow cytometric analysis showing the ratio of apoptotic (Annexin V^+^, PI^-^) and necrotic (Annexin V^+^, PI^+^) cells at 24 h and 48 h post addition of black, red, and green ink compared to the PBS controlled group. n=8 for 24h group. n=6 for 48 h group. Data are presented as mean ± SD. One-way **(A)** or Two-way **(F)** ANOVA followed by Bonferroni correction for multiple comparisons (* p < 0.05, ** p < 0.01, *** p < 0.001 **** p<0.0001).

### Tattoo ink induces an acute and long-lasting inflammation in the draining LN

Considering the relationship between the induction of cell death pathways and inflammation^27^, we hypothesized that ink-induced macrophage cell death could be associated with inflammation in the draining LN. Therefore, to evaluate the swelling of the LN, we estimated the total number of cells after tattooing with black ink, observing a significant increase at 48 h p.t., that followed a continuous growth for at least the first 240 h p.t (**Fig. 4A**). We observed similar dynamics in the number of CD45^+^ cells recruited to the pLN when the tattoo was made with the red and the green inks (**Supp. Fig. 4A**). The significant increase in the number of cells at 48 h p.t. was further confirmed by FACS analysis for B, T, NK cells, and conventional DCs, following tattooing with black ink (**Fig. 4B**), as well as with red and green inks (**Supp. Fig. 4B** and **4C**, respectively). To further characterize the inflammatory response, we collected the lymph from the pLN during the first 240 h following tattooing with black ink and we measured the levels of inflammatory cytokines and chemokines at different instances. According to the trend of the expression observed, we could identify an acute phase, characterized by the upregulation of the cytokines IL-6, KC, MCP-1, and MIP-1α, which peak at 6-12 h p.t., and return to basal levels after the first 240 h p.t. (**Fig. 4C**, upper graphs).

**Fig. 4.**
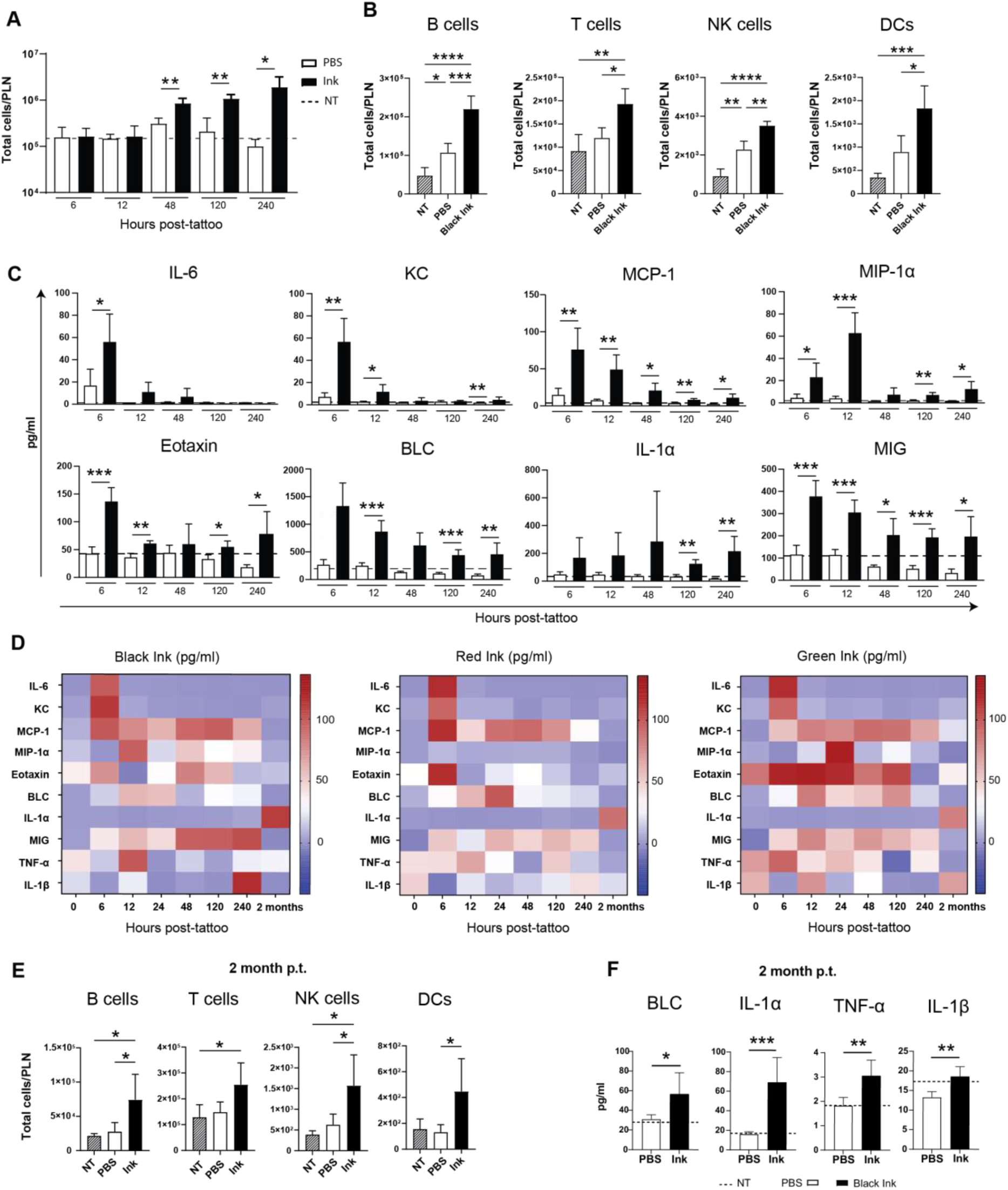
Characterization of the inflammatory response induced by tattoo ink in the lymphatic compartment. (**A**) Time course showing the total number of cells in the pLN during the first 240 h p.t. (**B**) Total number of B, T, NK cells, and DCs in the p LN at 48 h p.t. (**C**) Cytokine and chemokine levels in the draining LN during the 240 h that follows tattooing with black ink. Heatmap showing the expression of different cytokines and chemokines in the blood at various time points following tattooing with black (left panel), red (middle panel), and green ink (right panel). (**E**) Total number of B, T, NK cells, and DCs in the pLN at two months p.t. (**F**) Expression of BLC, IL-1α, TNF-α, and IL-1β, in the pLN at two months following black ink tattooing. In **A**-**F,** n=4 per group was representative of two independent experiments. Data are presented as mean ± SD. One-way **(B, E)** or Two-way ANOVA **(A, C)** followed by Bonferroni correction for multiple comparisons (* p < 0.05, ** p < 0.01, *** p < 0.001 **** p<0.0001).

Furthermore, we identified a second group of molecules, including eotaxin, BLC, IL-1α, and MIG, characterized by their long-term expression, which remains significantly elevated during the first 240 h p.t. (**Fig. 4C**, lower graphs). Additionally, we observed similar expression patterns when tattooing with both red and green inks (**Supp. Fig. 4D)**. To determine whether the observed inflammation also occurs systemically, we measured the expression of inflammatory mediators in blood at early (first five days) and late (two months) times post tattooing. Interestingly, we found that the early peak of inflammation previously described was also detected systemically, with elevated levels of the inflammatory cytokines IL-6, TNF-α and IL-1β, and the chemokines MCP-1, MIG and BLC, measured in the blood during the first 24 h following tattooing with all three tested inks (**Fig**. **4D**). The levels of most of these molecules were only transiently elevated. However, the levels of the alarmin IL-1α remain elevated in the blood from all tattooed groups two months after tattooing (**Fig. 4D**). Further, to characterize the long-term inflammatory response in the draining LN, we measured the numbers of B, T, NK cells, and DCs at two months p.t. with black ink (**Fig. 4D**), observing a significant increase in all the cell types at this time point, compared with the control groups (**Fig**. **4E**). We could also observe a tendency to have elevated levels of these cells in animals tattooed with red and green inks (**Supp. Fig. 4E and F**, respectively). Furthermore, we could also detect significantly elevated levels of the inflammatory cytokines BLC, IL-1α, TNF-α and IL-1β with black ink tattooing compared to PBS-tattooed control groups (**Fig. 4F**). Finally, to evaluate if the presence of the ink might be associated with higher cell proliferation in the draining LN, we performed immunohistology staining with KI-67, which demonstrated areas of cell proliferation in a LN associated with the area where ink accumulates, compared to areas without pigment (**Sup.** Fig. 4G).

### Tattoo ink impairs the immune response to a COVID-19 vaccine

To study the short - and long-term effects of tattooing on the antibody response following vaccination, we administered the Pfizer-BioNTech COVID-19 vaccine to mice at two different time points (two days or two months) after being tattooed with black, red, or green ink, and we evaluated the Ig responses against the receptor binding domain (RBD) of the spike protein. We found that all tattooed groups showed a significant decrease in the levels of anti-RBD specific IgG at ten days post-vaccination (p.v.) (**Fig. 5A**). In contrast, only the animals tattooed with red and green ink showed significantly decreased levels of IgM at seven days p.v. (**Supp. Fig. 5A**). Notably, the deficiency in the anti-RBD specific IgG responses was maintained in all the groups that were tattooed two months before. (**Fig. 5C**). However, we could not observe significant differences in the IgM response (**Supp. Fig. 5B**). To evaluate if the presence of the ink could affect the expression of the coronavirus spike protein in the medullary macrophages (MM), we performed immunostaining with an anti-spike antibody followed by FACS analysis at 24 h p.v. in two groups of mice tattooed 48 h or two months before. We observed that MM from all tattooed groups showed a tendency to display reduced levels of expression of the spike protein in the short-term experiments (**Fig. 5B**). Furthermore, at two months p.t., the expression levels of the spike protein were significantly decreased in MM from all the tattooed groups (**Fig. 5D**), compared to the control groups. Additionally, we observed a significant decrease in the expression of the costimulatory markers CD86 and CD80 in MM at 48 h and two months p.t. Specifically, CD86 levels were reduced in MM across all tattooed groups at 48 h p.t. (**Fig. 5E-F left panels**), while CD80 levels decreased in MM of the black and green ink tattooed groups at two months p.t. (**Fig. 5E-F middle panels**). Conversely, even if the costimulatory marker MHC-II tended to increase in all tattooed groups at 48 h p.t., the level declined by two months p.t (**Fig. 5E-F right panels**).

**Fig. 5.**
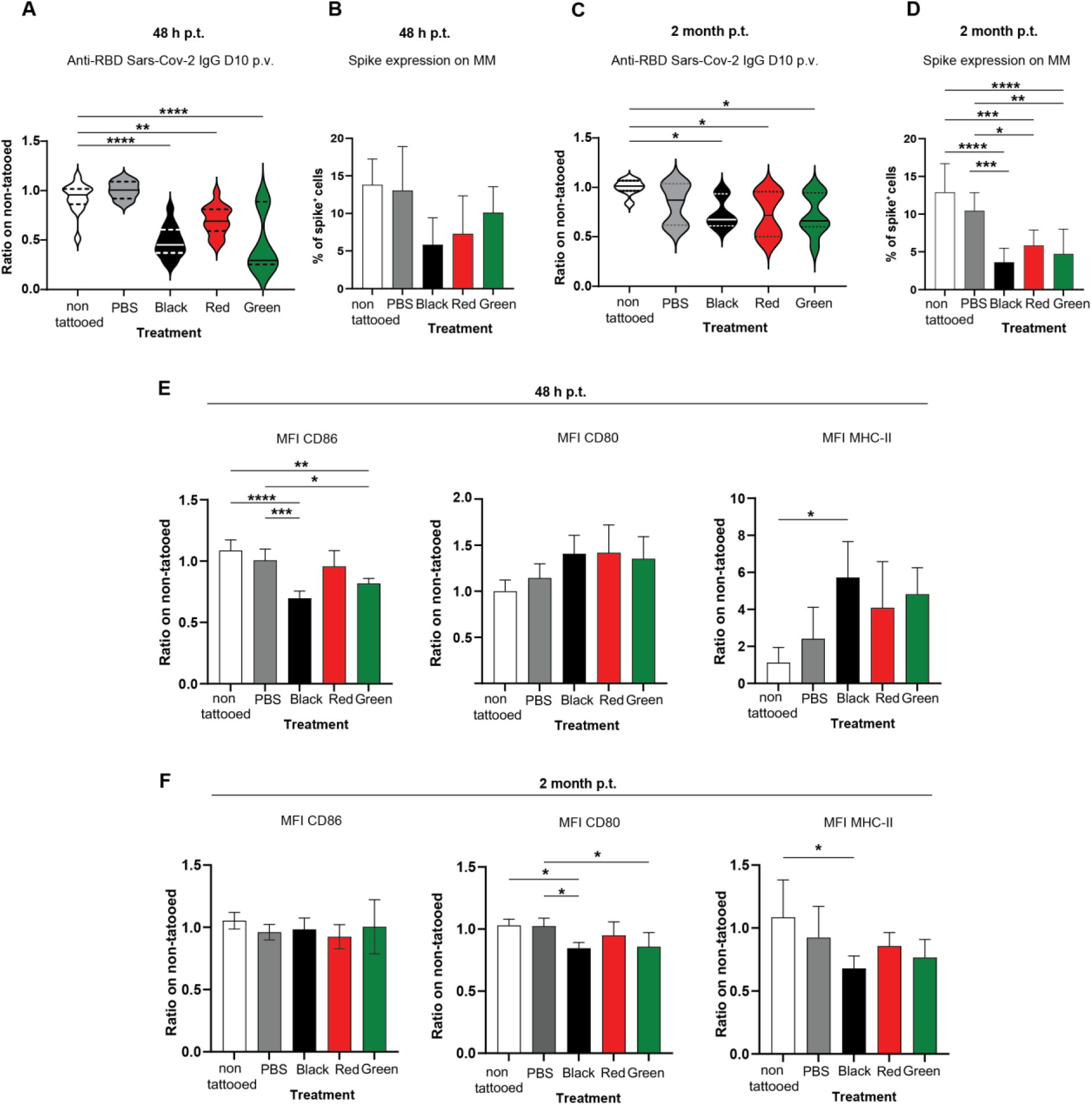
Characterization of the immune response to a COVID-19 vaccine after short and long-term tattooing. (**A**) Violin plot showing the anti-RDB Sars-CoV-2 IgG titters at day ten p.v. in animals tattooed 48 h before. n=14 for non-tattooed group, n=5 for PBS group, n=15 for black and red groups, and n=7 for green group. (**B**) Expression of coronavirus spike protein on medullary macrophages (MM) at 24 h p.v. in animals tattooed 48 h before. n=5 for 48 h p.t. group. n= 6 for 2 months p.t. group. (**C**) Violin plot of the anti-RDB Sars-CoV-2 IgG titters at day ten p.v. in long-term (two months) tattooed animals. n=8 for each group. (**D**) Expression of coronavirus spike protein at day ten p.v. in MM in a two-month post tattooing (p.t.) group. n=5 for 48 h p.t. group. n= 6 for 2 months p.t. group (**E**) Quantification of the expression of the costimulatory molecules CD86, CD80, and MHC-II in MM at 24 h p.v. in animals tattooed 48 h or two months (**F**) before vaccination. n=3 for non-tattooed group. n=6 for PBS group. n=7 for black, red, and green groups. Data are presented as median and 25th (bottom) and 75th (top) percentiles or mean ± SD. One-way ANOVA followed by Bonferroni correction for multiple comparisons (* p < 0.05, ** p < 0.01, *** p < 0.001).

To evaluate if the presence of ink could also affect the immune response in human cells, we differentiated macrophages from PBMCs of six healthy donors, added different amounts of the three tested inks together with the Pfizer-BioNTech COVID-19 vaccine, and co-culture them with B and T cells from the same donors. Interestingly, we observed that the presence of all inks significantly reduced the anti-spike specific IgG produced by B cells at day nine p.v. with the tested concentrations of ink (**Supp. Fig. 5C**). However, as in the case of the mouse, the levels of IgM produced were only significantly reduced after treatment with the red and green inks (**Supp. Fig. 5D**). Furthermore, as demonstrated in the murine model, the presence of black ink in the human macrophages significantly reduced the capture of the labeled vaccine (**Supp. Fig. 5E**), as well as the expression of the spike protein in these cells (**Supp. Fig. 5F**).

## Discussion

One of the urgent concerns associated with the safety of tattoos regards the potential redistribution of the unretained ink from the tattoo site to organs other than the skin and the toxic effect that the accumulation of these insoluble pigments might have at systemic levels. Previous studies have reported the deposition of pigments in the draining lymph node (dLN)^28–37^. Nevertheless, the contribution that the lymphatic vessels and bloodstream might play in this process needs to be further studied. Our work suggests that most unretained ink from the skin disseminates via the lymphatic system and accumulates in the medullary part of the dLN in the initial instances following tattooing. Moreover, we reported a progressive increase of ink pigment observed in the dLN two months post-tattoo, probably associated with constant draining from the tattooed site at the skin. This is particularly relevant considering that the puncturing of dermal blood vessels during the tattooing process in humans might also contribute to disseminating the ink via the bloodstream. Indeed, some studies have reported the presence of tattoo ink systemically associated with Kupfer cells of the liver, indicating that a blood-borne distribution of tattoo ink might also occur^40^. In addition, the extensive tattooed areas pose an added risk for the systemic distribution of the ink. Previous studies have estimated that, on average, 2.5 mg/cm^2^ are introduced into the dermal layer during the tattooing process^33,41^. Therefore, further studies are needed to evaluate, especially in people who get large tattoos, the potential correlation between the size of the tattoo and the accumulation of ink in internal organs, such as the spleen, liver, and kidney, as well as its pathophysiology consequences.

Although the adverse effects of tattoo pigment on the skin have been previously reported^42^, the impact of the accumulated pigments on the immune response is still unknown. In this work, we have demonstrated that macrophages exposed to different ink concentrations undergo apoptotic cell death with all the tested inks. A previous study has pointed out that monocytes and macrophages are the main immune cells involved in ink particle uptake after tattooing. It has been demonstrated that monocytes undergo significant alterations during their differentiation into macrophages, which could influence their viability and their sensitivity to tattoo ink^26^. The interaction of ink pigment with macrophages depends on their physicochemical characteristics and can lead to cytotoxic events, including cell viability perturbation via apoptosis or necrosis, oxidative stress, and reactive oxygen species (ROS) generation^43,44^. Furthermore, we have demonstrated that the ink pigments retained in the medullary region of the dLN stay associated with the phagocytic populations for months following tattooing. This is in line with other human studies in which ink pigments in dLNs were detected years after tattooing^47^. LN macrophages constantly filter the lymph and are essential for initiating innate immune responses that lead to the capture and inactivation of lymph-borne pathogens^48–51^. The accumulation of ink pigments, especially in the medullary region, is particularly relevant due to the highly phagocytic ability of these cells, specialized in the clearance of particulates, pathogens, and dying cells from the lymph and in support of plasma cell survival^52^. Considering the critical role of these cells in antimicrobial immunity, we hypothesize that tattoo ink pigment accumulation and the observed induction of macrophage death by the different types of ink could affect the capacity of these cells to control the spreading of pathogenic viruses and bacteria via the lymph, increasing the risk of dissemination. In this direction, previous studies have demonstrated that LN macrophages play a critical role in capturing lymph-borne viruses such as vesicular stomatitis virus (VSV), adenovirus, vaccinia virus, and murine cytomegalovirus^53^. Additionally, the depletion of LN macrophages before the VSV challenge led to a significant reduction in survival and an increase in viral titers found in the brain and spinal cord of the depleted mice^54^. The effect of the presence of ink in these cells on their antimicrobial function needs to be further evaluated in future studies.

The activation of the innate immune compartment is most likely associated with the prominent inflammatory response described in this study. This reaction in the lymphatic compartment could also be linked with the well-characterized acute inflammation caused by needles in the skin^55^, which might increase the chances of a dysregulated inflammatory response in patients who have developed allergic reactions or autoimmune diseases. Despite a common inflammatory trend observed in all tested pigments, significant differences were also associated with the capacity of certain pigments to induce inflammatory mediators. The initial acute inflammatory response was probably influenced, both at the cellular and molecular level, not only by the pigment alone but also by the presence of other ink components, such as binders, solvents, and additives known to be toxic^4^. These differences stressed the need for a careful evaluation of the safety profile of each of the chemical components of the tattoo ink mixture and the implementation of regulations that can reduce the heterogeneity in inks produced by different companies.

Additionally, different studies have previously associated chronic inflammation with multiple pathologies, including cancer^56^. In this direction, a recent article has confirmed the association between tattoo exposure and an increased risk of malignant lymphoma^57^. Furthermore, the elevated levels of the alarmin IL-1α in the draining LN were maintained during the first two months following tattooing with black ink. Interestingly, we have previously associated the elevated levels of this cytokine with the initiation of melanoma metastasis in the LN^58^. In addition, the observed chronic inflammation could also affect the immune surveillance and response to therapy in some tattooed cancer patients who undergo immune treatment^59^. A recent study analysed the usage of checkpoint BRAF/MEK inhibitors in tattooed patients with melanoma and the consequent adverse granulomatous reaction in the skin, which may be due to the enhanced loss of tolerance to tattoo inks^60^. Finally, the chronic inflammatory setting could potentially increase the carcinogenicity associated with certain pigments or their byproducts, increasing the risk of developing neoplasia^61^. Further studies are needed to identify the molecular basis of the connection between tattooing and cancer.

During the COVID-19 pandemic, one of the general concerns of the practitioners was the influence that administering a vaccine to a tattooed individual might have in developing the antibody response against the virus^62^. To date, no epidemiological studies have evaluated how the presence of a tattoo may affect the COVID-19 vaccine efficiency. In this work, we demonstrated, for the first time, the negative effect that the accumulation of ink has on the expression of the Spike protein in different antigen-presenting cells and, ultimately, in the humoral immune response elicited by the vaccine. Our results need to be further validated in a human study. However, different parameters need to be considered, such as the size and position of the tattoo and the type of ink included in the subject individuals of the study. Despite being more toxic, we could see an evident alteration of the immune response to vaccines in the animals tattooed with black ink, which might be associated with the capacity of carbon black to interfere with the antigen processing mechanism in the lymphocytic populations. Although we confirmed the detrimental role of tattoo ink in the response to the Pfizer COVID-19 vaccine, we cannot generalize these results to other types of vaccines, in which the mechanism of action might differ. Indeed, further studies assessing the effect of tattooing on the immune response induced by different vaccine types, including inactivated and attenuated vaccines and for several diseases, are needed to elucidate whether the effect found with the mRNA COVID-19 vaccine can be generalized. Conversely, tattooing might be an added risk for those individuals vaccinated with attenuated vaccines. Some reports associated with the live smallpox vaccination program in military service members have reported smallpox complications in different vaccinated, tattooed patients^67^. Future studies are needed to characterize these responses in more detail.

The toxicology risk of the components formed by tattoo inks remains a matter of concern^63^. A tattoo ink consists of a multi-component mixture of solvents, preservatives, additives, and non-liquid components like insoluble solid-state pigment particles. These pigments are responsible for the color of the tattoo. However, being a complex mixture of chemicals, different ink colors and suppliers contain different ingredients and impurities that may be responsible for the specific and heterogeneous cellular toxicity described. Further analyses are needed to investigate the mechanisms of transportation of such a complex mixture through the lymphatic system, analyzing which chemical components are transported with lymphatic fluid and lead, together with insoluble pigments, to the detected effects. The lack of regulation of the chemical component of the tattoo ink mixture is critical for this kind of analysis in which inks from different suppliers show a specific blend of ingredients with varying toxicity profiles. This study evaluated the three most used commercial tattoo inks: black (50%), red (14%), and green (9.1%)^64^. The observed differences in the toxicity effect of the tested inks are consistent with previous studies that identify red dyes as the most problematic, often associated with a cutaneous inflammatory response^65,66^. Red and black pigments are characterized by an aggregation of particles over time, resulting in larger pigment foreign bodies. This aggregation of pigments can be critical in developing local and systemic tattoo complications and can affect cellular uptake and transportation via the lymphatic system^23^.

In summary, this work represents the most extensive study to date regarding the effect of tattoo ink on the immune response and raises serious health concerns associated with the tattooing practice. Our work underscores the need for further research to inform public health policies and regulatory frameworks regarding the safety of tattoo inks.

## Material and Methods

### Mice

C57BL/6J mice were purchased from Charles River Laboratories. B6.129P2(Cg)-Cx3cr1tm1Litt/J (CX3CR1-GFP), B6.Cg-Tg(Itgax-Venus)1Mnz/J (CD11c-YFP) mice were initially purchased from Jackson Laboratories and subsequently bred in our animal facility at the Institute of Research in Biomedicine (IRB). All animal experiments were conducted following the guidelines of the Swiss Federal Veterinary Service, and the protocols (animal permits TI058/2021) were approved by the Animal Welfare Committee (Commissione Cantonale per gli Esperimenti sugli Animali) of the Cantonal Veterinary Office. Mice used for experiments included equal numbers of males and females, aged 6 to 12 weeks at the time of tattooing, in good health, and with no abnormal clinical signs. For the immunization study, ten μl of BNT162b2 mRNA vaccine (2μg per mouse) was injected into each footpad of isoflurane-anesthetized mice (2% volume-to-volume, Baxter), and pLN was collected at different time points. All mice in this study were bred under Specific Pathogen-Free (SPF) conditions in individually ventilated cages, with a controlled light-dark cycle (12:12), room temperature (20–24°C), and relative humidity (30%–70%). At the same time, all the experiments were run in Basics of Biosafety Level 2 (BSL2) facility.

### Tattoo model

Mice were anesthetized with a cocktail of ketamine (100 mg/kg, Sintetica) and xylazine (10 mg/kg, Virbac), injected intraperitoneally (IP), and immobilized. After disinfection of the footpad skin, an ATS-3 General Rodent Tattoo System (AIMS^TM^) machine was used to tattoo a small square area of the footpad, in a range between 20mm^2^ and 25mm^2^. The tattoo procedure was performed following standard healthy and hygienic recommendations. These include disinfection of the tattooed skin and aseptic oil (AIMS Tissue Oil / Skin Prep Cat#401, NSP01) to protect the skin during the first hours after the procedure, accelerating initial skin healing and preventing infection. Mice were randomly divided into experimental groups based on tattoo ink color (Intenze): Gen-Z Pure Green (Cat# ST1504GZP), Gen-Z True Black (Cat# ST1019TB), Gen-Z Bright Red (Cat #ST1007BR), the control group was tattooed with Phosphate-buffered saline without calcium and magnesium (PBS^-/-^). All tattooed mice were observed the following days for any discomfort measured by difficulty in walking, and the tattooed area was cleaned daily with a cotton stick soaked with an aseptic solution and oil during the healing period (max 15 days). Based on the experimental condition, mice were euthanized at specific time points, and pLNs or lumbar lymph nodes were collected.

### Chemical analysis of inks

Tattoo inks may contain undisclosed components; therefore, the identity of the pigments in the red and green ink was checked with RP-HPLC-DAD after extraction of the pigments with N and N-Dimethylformamide. All three inks were analyzed for educts and impurities from synthesizing the pigments and other undisclosed ingredients with RP-UHPLC-DAD. Polyaromatic hydrocarbons (PAH) were investigated in black ink using HPLC with fluorescence detection (FLD) after microwave extraction at 120°C with toluene. Form- and acetaldehyde, carcinogenic impurities often found in tattoo inks, were investigated with RP-HPLC-DAD after inline derivatization with 2,4-dinitrophenylhydrazine. Further, all inks were checked for more than 50 preservatives. Samples were extracted with methanol containing 0,1% phosphoric acid and then analyzed with RP-UHPLC-DAD.

### Two-Photon Intravital Microscopy (2PIVM)

Mice were anesthetized with a cocktail of ketamine and xylazine, as described in the section on the tattoo model. The mouse was positioned on a microscopic stage. The right foot was maintained and extended by fastening the fingers using a suture thread, and fasteners were attached to the hip. Finally, the lymphatic and haematic vessels were exposed through an incision in the skin (around 5 mm) and by removing the overlying fat tissue. Lymphatic vessels were labeled by subcutaneous injection into the right footpad of 70 kDa Rhodamine B isothiocyanate-Dextran. Mice were then tattooed with Gen-Z Bright Red ink, as previously described. The imaging of lymphatic and haematic transport of the ink was performed on a customized up-right two-photon platform (TrimScope, LaVision BioTec) as previously described^68^. The objective used was a 10X and two-photon micrographs were acquired every 5 min for a total duration of 60 min. Analyses were performed using the Imaris 9.9.1 software (Oxford Instruments) to obtain the 3D rendering of the vessels.

### Immunofluorescence

Popliteal LNs were harvested and fixed in 4% paraformaldehyde (PFA; Merck-Millipore) for 12 h at 4° and then embedded in 4% Low Gelling Temperature Agarose (Sigma-Aldrich). Sequential sections of 50 μm were cut with a vibratome (VT1200S, Leica Microsystems) and stained in a blocking buffer composed of PBS supplemented with calcium and magnesium (PBS^+/+^), TritonX100 (VWR) 0.1%, BSA 5% (VWR), αCD16/32 diluted 1:100. Slices were stained for: CD21/35 Pacific Blue, F4/80 AF488, CD11c AF594, CD169 AF647. After o/n incubation at 4 °C, samples were washed in PBS^+/+^ with 0.05% Tween 20 (Sigma-Aldrich), followed by washes with PBS^-/-^, and mounted on glass slides. Immunofluorescence confocal images were acquired using a Leica TCS SP5 confocal microscope (Leica Microsystems). Images were processed using Fiji and Imaris software (Oxford Instruments, v9.9.1). An additional imaging channel, specific for the ink, was generated by classifying each pixel as foreground or background. pLN regions were manually identified based on CD169 and CD21/35 expression. The colocalization analyses of the ink channel and specific cell staining were achieved using the Coloc functionality of Imaris.

### Electron microscopy

For electron microscopy, organs were immediately fixed after collection in 2% formaldehyde, 2,5% glutaraldehyde, and 2 mM CaCl_2_ in 0,15M cacodylate buffer pH 7.4 for 12 h at 4°C. Samples were then postfixed sequentially with 2% OsO4 in 0,1M cacodylate for three hours and 2,5% potassium ferricyanide in 0,1M cacodylate for 90 min at room temperature. After several washes in MilliQ, water samples were incubated overnight at 4 C° in 0,5% uranyl acetate. Samples were progressively dehydrated in ethanol, rinsed in propylene oxide, and infiltrated in a mixture of propylene oxide/epoxy resin (Epoxy Embedding Medium kit, Sigma-Aldrich) overnight at room temperature. Samples were finally embedded in pure resin and baked at 60°C for 48 h.

Semithin sections (1μm) were cut using a Leica EM UC7 ultramicrotome (Leica Microsystems), collected on superfrost microscope slides (Thermo Scientific), and stained with toluidine blue (1% toluidine in 1% sodium borate). Slides were acquired with an Aperio AT2 Slide scanner using Imagescope software (Leica Biosystems). Ultrathin sections (50-70nm) were collected using a Leica EM UC7 ultramicrotome (Leica Microsystems) on carbon-coated formvar slot grids and analyzed with a TALOS L120C transmission electron microscope (Thermo Fisher Scientific). Images were captured with a CETA 16M camera using Velox or MAPS software (Thermo Fisher Scientific).

### Hematoxylin and Eosin (H&E) Staining and Immunohistochemistry

Lymph nodes were fixed in 4% buffered formalin, embedded in paraffin, cut at 4 µm of thickness and stained with hematoxylin and eosin (HE). Immunohistochemical staining was performed on a Ventana Benchmark Ultra automated staining platform using a CC1 antigen retrieval buffer (Roche-Ventana) for 30 minutes at 100 C. This was followed by incubation with a monoclonal mouse anti-human CD68 antibody (clone Kp-1, Cell Marque, dilution 1:400; Biosystems, Switzerland) or with a monoclonal mouse anti-human CD163 antibody (MRQ26, Roche, ready to use) for 32 min., followed by the Optiview HRP multimer secondary detection system (Roche Ventana).

### Cell culture

Bone marrow-derived macrophages (BMDMs) were differentiated from monocyte precursors as previously described^69^. Briefly, bone marrow cells were isolated from femurs of 8 to 12-week-old WT C57/Bl6 mice and plated in complete DMEM (Gibco) without serum o.n. at 37°C with 5% CO_2_. Nonadherent cells were recovered and plated at 0.5/0.8 × 10^6^cells/ml and cultured for 7 days in complete DMEM supplemented with 10% fetal bovine serum (FBS; Gibco), 100 U/mL penicillin/streptomycin (Gibco), 2 mM L-glutamine (Gibco) and 5 ng/mL murine macrophage-colony-stimulating factor (mM-CSF; PeproTech). Human macrophages were differentiated from monocytes as previously described^70^. Briefly, monocytes were obtained from healthy blood donor buffy coats by two-step density gradient centrifugations. Peripheral blood mononuclear cells (PBMCs) from healthy volunteers were isolated as previously described^71^ using Ficoll Paque Plus (Sigma Aldrich Chemie) and 46% Percoll (VWR International) followed by incubation of purified cells in RPMI 1640 (Lonza) without serum, for 6 h at 37°C with 5% CO_2_. Adherent monocytes were cultured for 7 days in a complete RPMI medium (Gibco) supplemented with human M-CSF (50 ng/ml; PeproTech) to obtain completed differentiated monocytes-derived macrophages (MDMs). After 7 days of differentiation, both BMDMs and MDMs were detached with 0.05M EDTA (PanReac Applichem) in PBS^-/-^ at 37°C for 15 minutes with 5% CO_2_ and plate at a concentration of 0. 5 × 10^6^ cells/ml in complete DMEM (Gibco) in 24 well plates. The day after cells were incubated with the three selected inks or PBS^-/-^ for one h, washed with complete DMEM, and treated based on the experimental conditions.

### Cytospin

MDMs and MDMs were cultured and treated with ink as described in the paragraph Cell Culture. 24 h post ink removal cells were detached with 0.05M EDTA (PanReac Applichem) in PBS^-/-^ at 37°C for 15 min, pelleted and resuspended in complete RPMI 40% FBS. 100 μl of cell suspension was charged into each Cytofunnel (Double Cytofunnel^TM^, Epredia) and centrifuged for 4 min at 1000 rpm. Non-polarized slides (Epredia^TM^) with cells were left at RT until the slide was completely dry and then stained with RAL Diff-Quik™ (Siemens Healthcare Diagnostics AG) following manufacturer instructions. Finally, slides were mounted with a quick passage in Xylenes (Sigma-Aldrich) and assembled with Eukitt® Quick-hardening mounting medium (Sigma-Aldrich).

### ImageXpress

BMDMs were cultured as described above. On day 7 cells were detached with 0.05M EDTA (PanReac Applichem) in PBS^-/-^ at 37°C for 15 min and seeded at 8 × 10^4^ cells/well in a µ-Slide 8 Well high Glass Bottom (Ibidi), previously coated with Poly-L-lysine solution 0.01% (m/v) in dH20 for 30 min at RT. After ON incubation at 37°C and 5% CO_2,_ BMDMs were treated for 1 h with selected inks at 37°C and 5% CO_2_. After ink removal, cells were washed with PBS^-/-^ and stained with Hoechst (Thermo Fisher Scientific) 1μg/ml and PI (Sigma-Aldrich) 2.5 μg/ml in complete DMEM without Phenol Red for 30 min at 37°C with 5% CO_2_ and imaged with an ImageXpress Micro High Content Screening System (Molecular Devices). To image plates and for data analyses, MetaXpress software was used with settings as follows: acquisition every 2 h, 20X objective; double site to visit; wavelengths of DAPI and Texas Red.

### Flow Cytometry

Flow cytometry analysis of cell death in BMDMs and MDMs. Macrophages were treated with inks as described in the paragraph Cell culture and, after 24 or 48 h, were detached with 0.05M EDTA (PanReac Applichem) in PBS^-/-^ at 37°C for 15 min. Fc receptors were blocked with αCD16/32, and BMDMs were detected using an αF4/80, while αCCR7 and αCCR2 were used for MDMs. Surface staining was carried out for 30 min RT in the dark. The APC Annexin V Apoptosis Detection Kit with 7-AAD (BioLegend) was used to identify apoptotic and necrotic cells following manufacture instructions. To analyze immune infiltrate in pLN, the LNs were collected, disrupted with tweezers, and filtered the resulting cells with a 40 µm cell strainer. A complete list of the mAbs used is reported in Table 2. Fc receptors were blocked with αCD16/32, and dead cell exclusion was performed using the Zombie Aqua^TM^ Fixable Viability Kit (Biolegend, San Diego, USA). Surface staining was carried out for 30 min, and MMs were detected as F4/80^+^/CD11b. All operations were done at 4°C in the dark. For the analysis of the capture of vaccines and spike expression by human macrophages after the incubation with the DiD-labelled BNT162b2 mRNA vaccine (Pfizer/BioNTech), as described in the paragraph “*In vitro* human co-culture of Macrophage and Lymphocyte with DiD-labelled BNT162b2 mRNA vaccine”, cells were harvested by incubating them with 100 µl of PBS^-/-^ with 0.05M EDTA for 15 min at 37°C with 5% CO_2_. After staining with a viability dye, cells were surface stained with an anti-spike antibody. All operations were done at 4°C in the dark. All samples were acquired on fluorescence-activated cell sorting (FACS) Symphony A3 flow cytometer (BD Biosciences, New Jersey, USA) and were analyzed with FlowJo software version 10.7.1 (FlowJo LLC, Ashland, Oregon).

### Multiplex assay

Blood was collected, at different time points post tattooing, from the tail vein in capillary blood collection system tubes (GK 150 SE Gel 200 µl, Kabe Labortechnik). Blood was kept for one h at RT to clot and then centrifuged at 3,000g for 10 min RT. The serum was aliquot into 1.5 ml Eppendorf and stored at -80°C until the usage. pLNs were collected at different time points after tattooing and carefully disrupted in 75μL of cold PBS^-/-^to avoid cell rupture. The suspension was centrifuged at 1,500 rpm for 5 min, and the supernatant was collected. The concentration of cytokines and chemokine in the serum and lymph was determined by LEGENDPlex assays (Mouse Proinflammatory Chemokine Panel and Mouse Inflammation Panel (13-plex); Biolegend) according to the manufacturer’s instructions. Samples were analyzed using a Symphony A3 flow cytometer (BD Biosciences, New Jersey, USA), and data were analyzed using LEGENDPlex software (BioLegend).

### Vaccine storage and labeling

Discarded remnant material from the BNT162b2 mRNA vaccine (Pfizer/BioNTech) was stored at 100 µg/ml at -80°C. The vaccine was incubated with 0.40 µM DID dye for 10 min at RT for labeling. Following incubation, the vaccine was centrifuged using Amicon® Ultra Centrifugal Filter (10 kDa MWCO) at 800g for 60 min.

### ELISA

SARS-CoV-2 RBD protein was produced in-house as previously described^72^. Blood was collected for *in vivo* immunological assays as described in the Multiplex assay paragraph on days seven and 10 p.v. Serum was isolated by centrifugation for 10 minutes at 3000g and stored at -80°C for later analysis. Half-well Nunc ELISA plates were coated with 5μg/ml SARS-CoV-2 RBD protein in PBS^-/-^ buffer and incubated overnight at 4°C. After blocking with PBS^-/-^ containing 1% of Bovine Serum Albumin (BSA), 25 μl of diluted serum samples were added to each well and incubated for 2 hours at RT. Alkaline phosphatase (ALP)-conjugated αIgG and αIgM antibodies (SouthernBiotech, 1:500 dilution) in PBS^-/-^ containing 1% of BSA were used to detect antibody binding. The Alkaline Phosphatase Yellow (pNPP) Liquid Substrate System for ELISA (Sigma-Aldrich) was used as a substrate, and the reaction was stopped by adding an equal volume of 0.3M NaOH. Absorbance was read on a GloMax® Discover Microplate Reader (Promega) at 405 nm.

For the *human in vitro co-culture* immunological assays, supernatant was collected on days 6 and 9 p.v., centrifuged for 5 minutes at 1500 rpm, and stored at -80°C for later analysis. The assay was performed as described above. Horseradish peroxidase (HRP)-conjugated αIgG and αIgM antibodies (SouthernBiotech, 1:5000 dilution) in PBS^-/-^ containing 1% of BSA diluted were used to detect antibody binding. TMB/E Ultra-Sensitive, Blue, Horseradish Peroxidase Substrate (Merk) was used as the substrate, and the reaction was stopped by adding an equal volume of 1M H_2_SO_4_. Absorbance was read on a GloMax® Discover Microplate Reader (Promega) at 450 nm.

### *In vitro* human co-culture of macrophages and lymphocytes with DiD-labelled BNT162b2 mRNA vaccine

Human monocyte-derived macrophages were differentiated and seeded by adding 0.1 × 10^6^ cells/ml per well. The cells were treated as described in the paragraph cell culture. They were then pulsed for 5 h with 1μg/ml DiD-labelled BNT162b2 mRNA vaccine (Pfizer/BioNTech), which had previously been incubated in complete RPMI for 60 min at 37°C with 5% CO_2_ to allow the formation of the protein corona. After incubation, the supernatant was removed and replaced with lymphocyte-containing medium.

From the lymphocyte fraction of the PBMCs, B, and T cells were isolated as described in the cell culture paragraph. Lymphocytes were thawed from cryo-storage, washed with the B-cell medium, and centrifuged at 1500 rpm for 5 min. They were stained with CellTrace Violet (Thermo Fisher Scientific) following manufacturer instructions. Cells were washed using complete RPMI, and 200 μl of 1.25 × 10^4^ cells/ml lymphocytes were co-cultured in duplicate with ink-treated macrophages. The cultures were treated with IL-21 enriched B-cell medium (50 ng/ml, Peprotech, London W6 8LL, UK) in 96-well flat-bottom plates for six or nine days at 37°C with 5% CO_2_.

## Acknowledgments

Swiss National Science Foundation grant 204636 (CP, LR,KC, SG), Swiss Centre for Applied Human Toxicology (SCAHT) (to AC) funding project 2022-2024, Swiss Government Excellence Scholarship (to JF), Fidinam Foundation.

## Authors contributions

Conceptualization: AC, JF, SG; WB, MP, DR; Experiments: AF, JF, SG, CP, IL, LM, VS, UH, AR, TV, SM, LG, LR, MF, KC; Data analysis and visualization: AC, JF, AP, CP, LM, VS, SG; Writing original draft: AC, JF, SG.

## Competing interest

The authors declare that they have no known competing financial interests or personal relationships that could have appeared to influence the work reported in this paper.

Where authors are identified as personnel of the International Agency for Research on Cancer/ or the World Health Organization, the authors alone are responsible for the views expressed in this article and the views do not necessarily represent the decisions, policy, or views of the International Agency for Research on Cancer/ or the World Health Organization.

Supplementary Information is available for this paper.

## Supplementary Tables

**Table 1.**
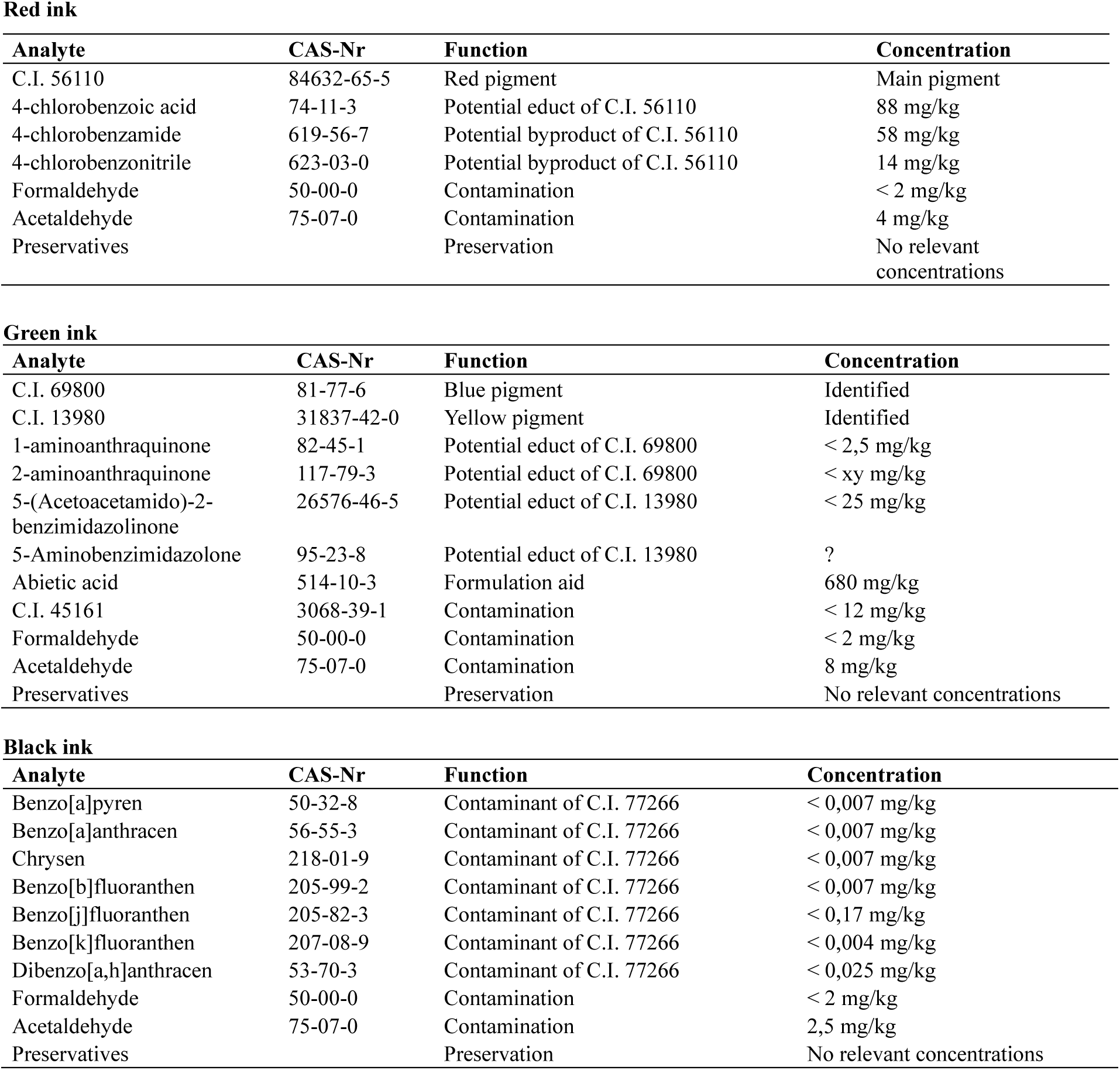
Chemical analysis data of the inks.

Red ink
| Analyte | CAS-Nr | Function | Concentration |
| --- | --- | --- | --- |
| C.I. 56110 | 84632-65-5 | Red pigment | Main pigment |
| 4-chlorobenzoic acid | 74-11-3 | Potential educt of C.I. 56110 | 88 mg/kg |
| 4-chlorobenzamide | 619-56-7 | Potential byproduct of C.I. 56110 | 58 mg/kg |
| 4-chlorobenzonitrile | 623-03-0 | Potential byproduct of C.I. 56110 | 14 mg/kg |
| Formaldehyde | 50-00-0 | Contamination | < 2 mg/kg |
| Acetaldehyde | 75-07-0 | Contamination | 4 mg/kg |
| Preservatives |  | Preservation | No relevant concentrations |

Green ink
| Analyte | CAS-Nr | Function | Concentration |
| --- | --- | --- | --- |
| C.I. 69800 | 81-77-6 | Blue pigment | Identified |
| C.I. 13980 | 31837-42-0 | Yellow pigment | Identified |
| 1-aminoanthraquinone | 82-45-1 | Potential educt of C.I. 69800 | < 2,5 mg/kg |
| 2-aminoanthraquinone | 117-79-3 | Potential educt of C.I. 69800 | < xy mg/kg |
| 5-(Acetoacetamido)-2-benzimidazolinone | 26576-46-5 | Potential educt of C.I. 13980 | < 25 mg/kg |
| 5-Aminobenzimidazolone | 95-23-8 | Potential educt of C.I. 13980 | ? |
| Abietic acid | 514-10-3 | Formulation aid | 680 mg/kg |
| C.I. 45161 | 3068-39-1 | Contamination | < 12 mg/kg |
| Formaldehyde | 50-00-0 | Contamination | < 2 mg/kg |
| Acetaldehyde | 75-07-0 | Contamination | 8 mg/kg |
| Preservatives |  | Preservation | No relevant concentrations |

Black ink
| Analyte | CAS-Nr | Function | Concentration |
| --- | --- | --- | --- |
| Benzo[a]pyren | 50-32-8 | Contaminant of C.I. 77266 | < 0,007 mg/kg |
| Benzo[a]anthracen | 56-55-3 | Contaminant of C.I. 77266 | < 0,007 mg/kg |
| Chrysen | 218-01-9 | Contaminant of C.I. 77266 | < 0,007 mg/kg |
| Benzo[b]fluoranthen | 205-99-2 | Contaminant of C.I. 77266 | < 0,007 mg/kg |
| Benzo[j]fluoranthen | 205-82-3 | Contaminant of C.I. 77266 | < 0,17 mg/kg |
| Benzo[k]fluoranthen | 207-08-9 | Contaminant of C.I. 77266 | < 0,004 mg/kg |
| Dibenzo[a,h]anthracen | 53-70-3 | Contaminant of C.I. 77266 | < 0,025 mg/kg |
| Formaldehyde | 50-00-0 | Contamination | < 2 mg/kg |
| Acetaldehyde | 75-07-0 | Contamination | 2,5 mg/kg |
| Preservatives |  | Preservation | No relevant concentrations |

**Table 2.** Antibody list.

| Antibody | Fluorophore | Company | Cat. Num. | Clone | lot |
| --- | --- | --- | --- | --- | --- |
| F4/80 | FITC | Biologend | 123120 | BM8 | B380534 |
| CD86 | PercCP | Biologend | 105026 | GL-1 | B343360 |
| Gr1 | APC-Cy7 | Biologend | 108424 | RB6-8C5 | B388923 |
| MHC II | PB | Biologend | 107620 | M5/114.15.2 | B385751 |
| Spike | BV750 | Biotechne | FAB105805S | 1035226 | 1731506 |
| CD80 | BV605 | Biologend | 104729 | 16-10A1 | B357122 |
| CD11c | BV711 | Biologend | 117349 | N418 | B377887 |
| CD11b | BUV395 | BD Bioscience | 563553 | M1/70 | 3346840 |
| B220 | BUV496 | BD Bioscience | 612950 | RA3-6B2 | 3226463 |
| CD45 | BUV563 | BD Bioscience | 612924 | 30-F11 | 4043831 |
| NK1.1 | BUV615 | BD Bioscience | 751111 | PK136 | 4093293 |
| CD169 | PE | Biologend | 142404 | 3D6.112 | B364410 |
| CD3 | PECY7 | Biologend | 100220 | 17A2 | B389532 |
| CD21/35 | Pacific Blue | Biologend | 123414 | 7 E9 | B316532 |
| CD11c | AF594 | Biologend | 117346 | N418 | B273696 |
| CD169 | AF647 | Biologend | 142408 | 3D6.112 | B350284 |
| CD16/32 | Purified | Biologend | 101302 | 93 | B401727 |
| Annexin V | APC | Biologend | 640920 |  | B397838 |
| 7-AAD Viability staining solution |  | Biologend | 420404 |  | B383019 |
| Zombie aqua fixable viability kit |  | Biologend | 423101 |  | B398272 |
| CCR2 | PE | Biologend | 357205 | K036C2 | B404134 |
| CCR7 | PE/cyanine 7 | Biologend | 353226 | G043H7 | B257474 |
| CD19 | BUV737 | BD Bioscience | 612756 | SJ25C1 | 3153506 |
| IgG | PE-CF594 | BD Bioscience | 562538 | G18-145 | 3282863 |
| CD4 | BUV496 | BD Bioscience | 750980 | OKT4 | 4117147 |
| CD38 | APC | Biologend | 303510 | HIT2 | B398666 |
| CD3 | APC/cyanine7 | Biologend | 300318 | HIT3a | B364230 |
| CD154 | BV605 | Biologend | 310826 | 24-31 | B392352 |
| CD138 | BV711 | Biologend | 356522 | MI15 | B379497 |
| CD27 | PE/cyanine 7 | Biologend | 356412 | M-T271 | B396581 |
| CD28 | PerCP/Cyanine 5.5 | Biologend | 302922 | CD28.2 | B384456 |

## Supplementary Figures

**Supp. Fig. 1.**
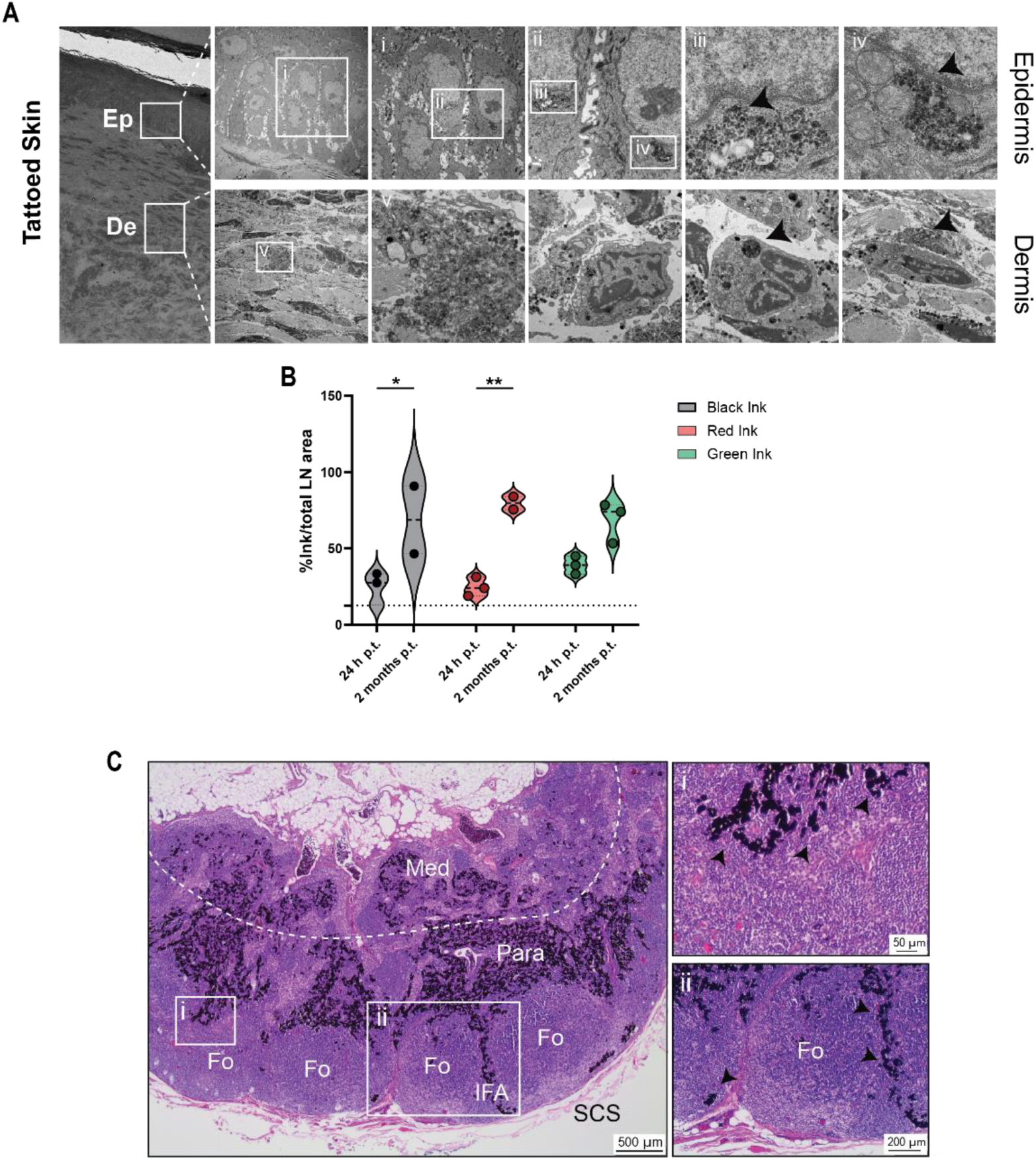
Tattoo ink not retained in the skin accumulates in the draining lymph node. **(A)**, Representative EM micrographs showing the presence of ink in the epidermis and the dermis of the footpad of a tattooed animal at 24 h p.t. The ink (black arrowhead) is associated with different types of cells, including keratinocytes (Epidermis, i-iv) **(B)** Violin plot representing the % of the area occupied by the ink on the total area of the draining LN at 24 h and two months p.t. Data are presented as median and 25th (bottom) and 75th (top) percentiles. One out of two or more independent experiments is shown Two-way ANOVA followed by Bonferroni correction for multiple comparisons (* p < 0.05, ** p < 0.01, *** p < 0.001 **** p<0.0001). **(C)** Haematoxylin-Eosin staining of the LN of a patient who presents a tattoo. The ink accumulates in the medullary (Med), paracortical (Para), and interfollicular area (IFA).

**Supp. Fig. 2.**
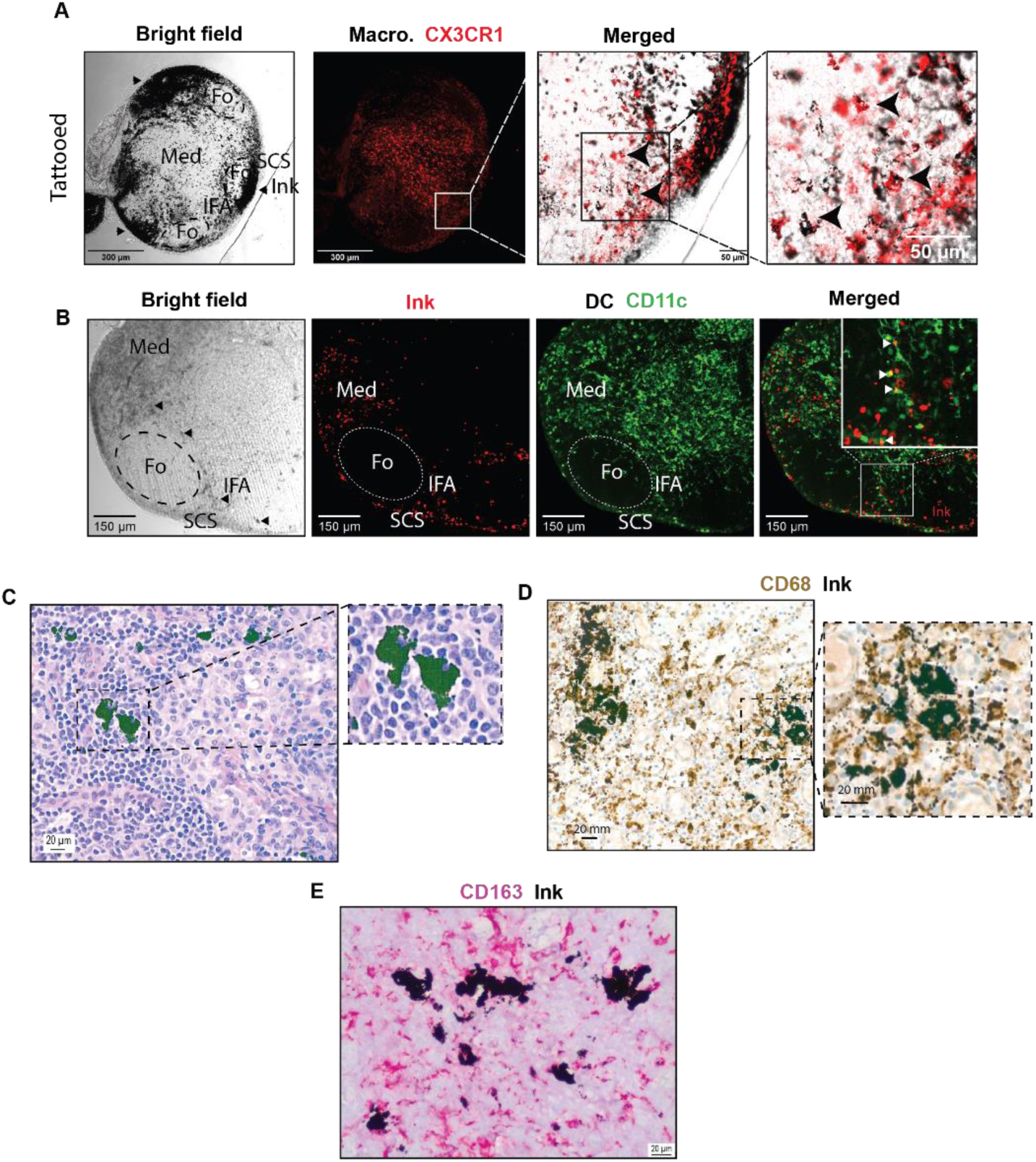
Medullary macrophages capture and retain tattoo ink. (**A**) Representative microscopy bright field and confocal images of tattooed GFP-expressing CX3CR1 mice. Black arrows indicate macrophages containing ink vesicles (right). (**B**) Representative correlative microscopy image of tattooed YFP-expressing CD11c mice. White arrows indicate DC associated with ink (right). (**C**) H&E staining of a human LN from a tattooed patient shows giant cells associated with tattoo pigment. (**D-E**) Representative immunohistochemical staining confirms that the giant cells associated with tattoo pigment express the macrophage markers CD68 and CD163.

**Supp. Fig. 3.**
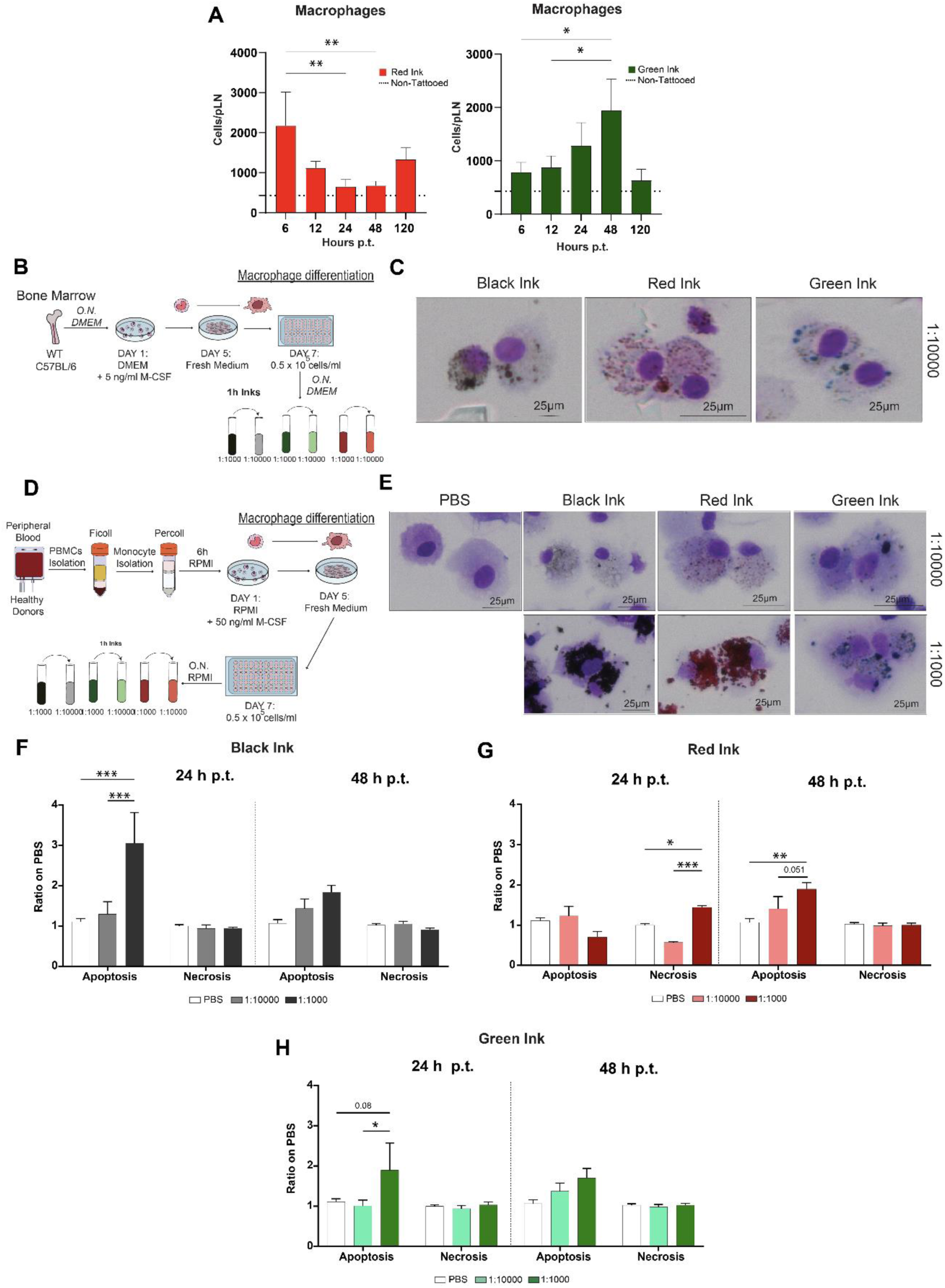
Tattoo ink induces macrophage death. **(A)**, Absolute number of macrophages in the pLN of tattooed animals during the first 120 h following footpad tattooing with red (left) and green (right) ink. n=4 representative of two independent experiments. (**B**) Schematic drawing of the protocol followed for the macrophage differentiation in the murine and human (**D**) settings. (**C**) Representative cytospin images showing ink uptake by murine-derived or human-derived (**E**) macrophages. (**F-H**) Flow cytometric analysis showing the ratio of apoptotic (Annexin V^+^, PI^-^) and necrotic (Annexin V^+^, PI^+^) cells at 24 h (left) and 48 h (right) post addition of black, red, and green ink, compared to PBS control. N=5. Data are presented as mean ± SD. One-way **(A)** or Two-way **(F-H)** ANOVA followed by Bonferroni correction for multiple comparisons (* p < 0.05, ** p < 0.01, *** p < 0.001 **** p<0.0001).

**Supp. Fig. 4.**
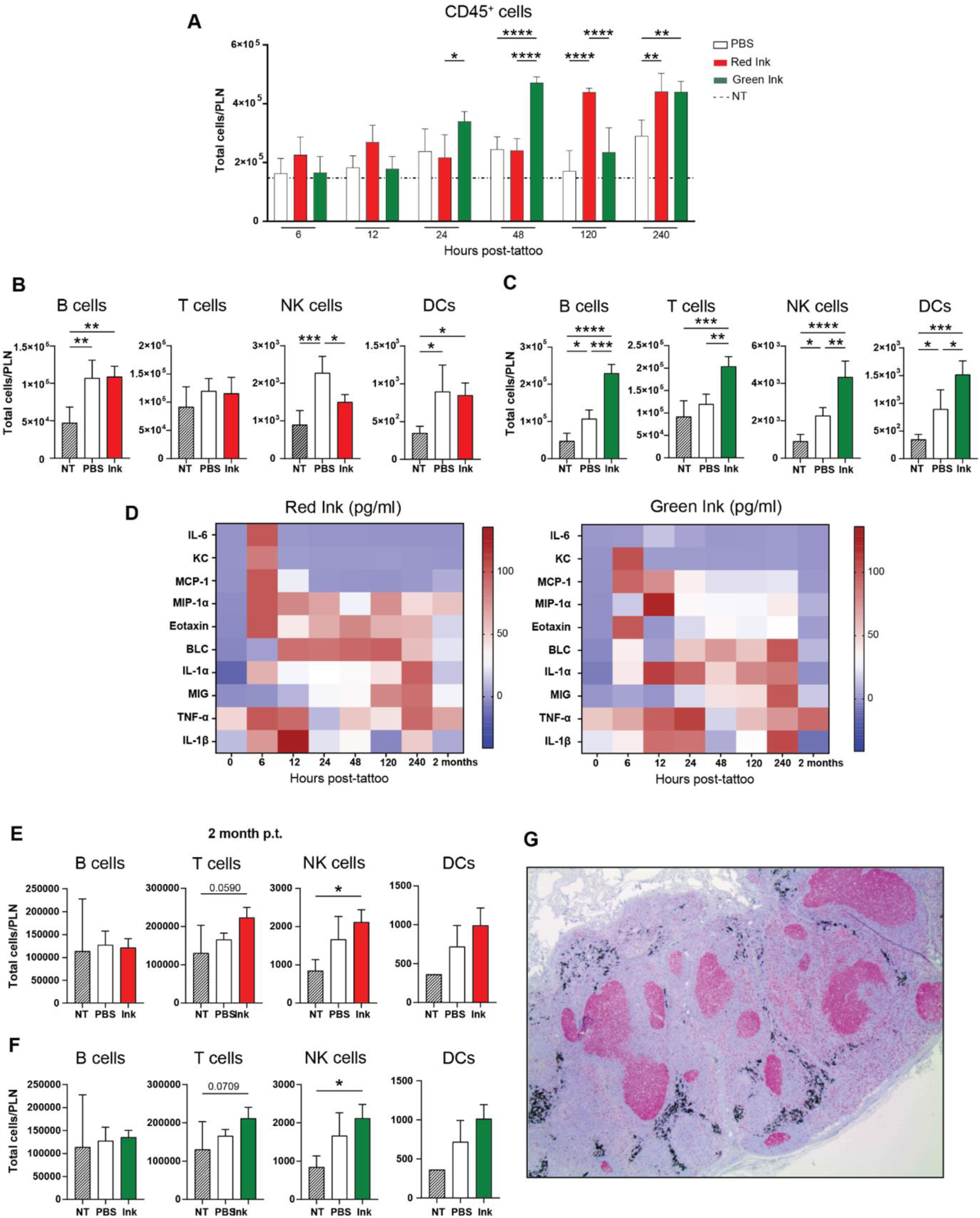
Characterization of the inflammatory response induced by tattoo ink in the lymphatic compartment. (**A**) Time course showing the total number of CD45^+^ cells in the pLN during the first 240 h following tattooing with red (red bars) and green (green bars) ink. NT stands for non-tattooed. Total number of B, T NK cells and DCs at 48 h and 2 months p.t with red ink (**B, E,** respectively) and green ink (**C, F,** respectively). (**D**) Heatmap showing the expression of different cytokines and chemokines in the lymph from the pLN at different time points following tattooing with red (left panel) and green (right panel) ink. (**G**) Immunohistochemical KI-67 staining of a human LN from a tattooed patient shows high proliferation areas in a LN associated with tattoo pigment. In **A**-**F** n=4 per group representative of two independent experiments. Data are presented as mean ± SD. One-way **(B, C, E, F)** or Two-way **(A)** ANOVA followed by Bonferroni correction for multiple comparisons (* p < 0.05, ** p < 0.01, *** p < 0.001 **** p<0.0001).

**Supplementary Figure 5.**
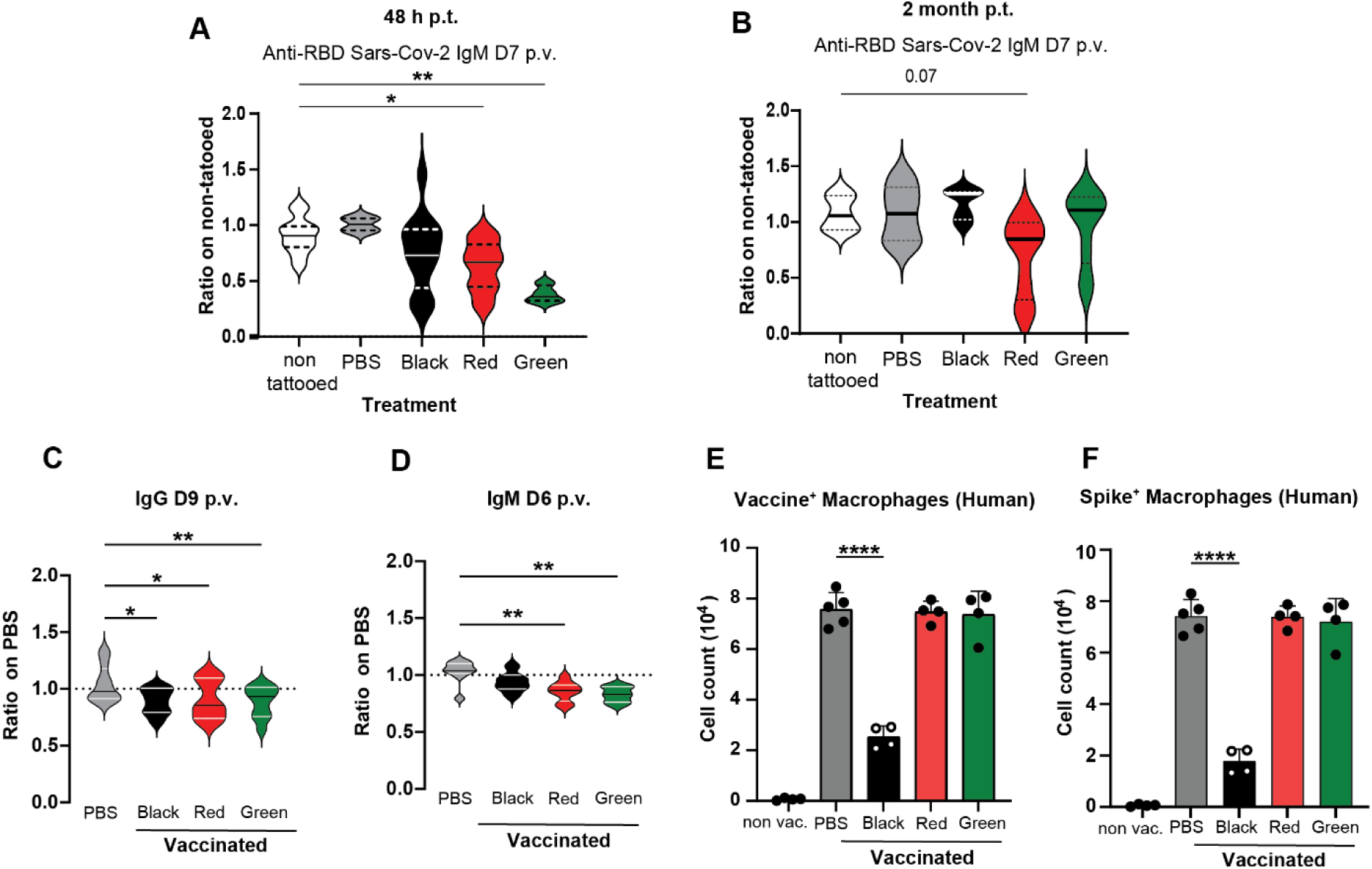
Tattoo ink impairs the immune response against a Covid vaccine. (**A**) Violin plots showed the anti-RDB Sars-CoV-2 IgM titters at day 7 p.v. in mice tattooed 48 h before, n=11 for the non-tattooed group, n=5 for PBS and green groups, n=13 for black and red groups. **(B)** Violin plots showed the anti-RDB Sars-CoV-2 IgM titters at day 7 p.v. in mice tattooed 2 months before, n=5 for the non-tattooed group, n=6 for the PBS group and n=8 for black, red, and green groups. (**C-D**) Violin plot of anti-RDB Sars-CoV-2 IgG at day nine p.v. and IgM at day six p.v. generated in vitro by co-culture of human PNMCs-derived macrophages with T and B cells from the same donors. n=9 for PBS group. n=6 for black, red, and green ink groups. (**E**) Capture of labeled coronavirus vaccine and expression of coronavirus spike protein (**F**) in human macrophages in vitro at five h p.v and measured by FACS analysis. Data are presented as median and 25th (bottom) and 75th (top) percentiles or mean ± SD. One-way ANOVA followed by Bonferroni correction for multiple comparisons (* p < 0.05, ** p < 0.01, *** p < 0.001, **** p < 0.0001).

